# The chromatin remodeler LSH controls genome-wide cytosine hydroxymethylation

**DOI:** 10.1101/2020.03.10.983148

**Authors:** Maud de Dieuleveult, Martin Bizet, Laurence Colin, Emilie Calonne, Martin Bachman, Chao Li, Irina Stancheva, François Fuks, Rachel Deplus

## Abstract

TET proteins convert 5-methylcytosine (5mC) to 5-hydroxymethylcytosine (5hmC), leading to a dynamic epigenetic state of DNA that can influence transcription. While TET proteins have been associated with either epigenetic repression or activation complexes, the overall understanding of the molecular mechanisms involved in TET-mediated regulation of gene transcription still remains limited. Here, we show that TET proteins interact with lymphoid-specific helicase (LSH), a chromatin remodeling factor belonging to the SNF2 super family. Lsh knock-out leads to a significant reduction of 5-hydroxymethylation global level in mouse embryonic fibroblasts (MEFs) and in embryonic stem cells (ESC). Whole genome sequencing of 5hmC in wild-type versus Lsh knock-out MEFs and ESCs showed that in absence of Lsh, some regions of the genome gain 5hmC while others lose it, with not much effect on gene expression. We further show that 5hmC modifications upon Lsh loss is not a direct consequence of 5mC decrease, as differentially hydroxymethylated regions (DhMR) did not overlap with DMR (differentially methylated regions), underlying that these modifications occurred at different genomic loci. Altogether, our results suggest that LSH is a key regulator of 5hmC in both MEFs and ESC and that TET proteins rely on specific factors to establish genome-wide 5hmC patterns.

## INTRODUCTION

DNA modifications and chromatin organization play a major role in a variety of biological and molecular processes. The best known modification base in DNA is 5-methylcytosine (5mC) established by the DNA methyltransferases (DNMTs) (1, 2). DNA methylation contributes to numerous biological processes, including embryonic development, X-chromosome inactivation, genomic imprinting and chromosome stability (3). The discovery of TET proteins, 2-oxyglutarate and Fe(II)-dependent dioxygenases which are able to oxidize 5mC in 5-hydroxymethylcytosine, 5-formylcytosine (5fC) and 5-carboxylcytosine (5caC), has revolutionized our understanding of DNA demethylation process (4–6). In addition to being an intermediate in DNA demethylation, 5hmC is now established as an epigenetic mark and has been shown to be the mostly stable in the mammalian genome (7, 8). New readers and modifiers are thus rising interest in getting deeper into the roles and the dynamics of 5hmC (9). Despite substantial research describing the genomic landscape of 5hmC in different cell systems, our understanding of the TETs and 5hmC regulation remains limited.

Approaches to elucidate how TET proteins function included search for interacting partners and led to the identification of transcriptional factors /nuclear receptors (CXXC4, NANOG, PPARγ, PU.1, EBF1, PRDM14), chromatin-associated proteins involved in transcriptional activation (OGT and SET1/COMPASS complex) (10) or repression (SIN3A/HDACs, NURD) (11). Also, insights have been obtained regarding the regulation of TET activity and expression. Ascorbic acid (vitamin C) has been shown to significantly enhance TET hydroxymethylation activity, acting as a specific co-factor for these enzymes probably by reducing Fe(III) back to Fe(II) after enzymatic catalysis (12–15). Tet1 can be regulated by several transcription factors as NF-κB with an anticorrelation with immune response genes in various cancer cell lines (16). Other groups indicated that TET proteins can be post-transcriptionally regulated by microRNAs (miRNAs) (17–20). Altogether, these results shed lights on regulatory mechanisms (post-translational modifications, miRNA network, small molecules) that impact TET expression and/or activity.

Various cancers and transformed cells harbor disturbed epigenomes, as illustrated by global DNA hypomethylation and focal promoter hypermethylation (21, 22). Interestingly, TET2 and IDH1/IDH2 have been found to be mutated in different cancers, including myeloproliferative neoplasms (MPNs), myelodysplastic syndromes (MDS) and acute myeloid leukemia (AML) (23–25). The regulation of TET proteins expression, without implying specific mutations, has also been involved in the development of cancers, for instance TET2 expression was significantly reduced in melanoma, causing global hypo-hydroxymethylation and hypermethylation of genes involved in disease progression (26). Although decreased 5hmC seems to be a common feature of various cancers, the precise mechanism linking TET deregulation and aberrant methylation is not properly understood yet.

The SNF2-like helicase LSH (also known as HELLS, SMARCA6 or PASG), initially identified as a factor required for lymphoid cells proliferation, belongs to the SNF2 family of chromatin remodeling factors (27, 28). Chromatin remodelers play important biological functions, such as transcriptional control, DNA repair and genomic recombination (29). LSH was shown to be able to slide nucleosomes *in vitro* (30, 31). Deletion of *Lsh* in mice was reported to be perinatally lethal with normal development, except for a small birth weight (32). Connection between Lsh and DNA methylation is well established. Its mutation or deletion leads to a profound loss of global DNA methylation, especially at repetitive sequences (33, 34) but also leads to hypo-as well as hypermethylation at specific genes (35). Knockdown of *Lsh* in embryonal carcinoma cells and ES cells, despite incomplete CG methylation, *Lsh-/-* embryos show cellular differentiation (36). In addition to reduced DNA methylation levels, the chromatin of *Lsh-/-* MEFs displays alterations in histone modifications, such as H3K4me3, H3K27me3 and H3K9me3 (37, 38).

Mechanistically, LSH acts as a recruitment platform for DNMTs, as it has been shown to interact with DNMT3A and DNMT3B and indirectly with DNMT1 and also CDCA7 (31, 39, 40). LSH may facilitate DNMTs recruitment by remodeling chromatin at their target genes. More recently, the ATP function of LSH was shown to be critical for nucleosome density and for complete cytosine methylation at specific loci, thereby implying that chromatin remodeling mediated by LSH is critical for the establishment of DNA methylation patterns (41). However, it is still unclear whether LSH loss affects the maintenance of DNA methylation patterns in addition to playing a role in de novo methylation (36, 39, 42, 43), when during development these effects occur and if other factors (apart from DNMTs) are involved in this process. Recently, new work has been done to understand the crosstalk between the two roles of LSH on methylation and nucleosome positioning suggesting that LSH is needed at enhancers and repetitive sequences to regulate the chromatin accessibility (41, 44).

In this study, we showed that LSH and TETs interact. We found that Lsh disruption leads to a reduction in the global level of 5hmC in MEFs and ESCs, which are well studied models for both TET and LSH. Nonetheless, genome-wide 5hmC studies in wild-type and Lsh^-/-^ MEFs and ESCs revealed that thousands of genomic regions gain or lose 5hmC. In most cases no changes in gene expression could be detected. Our data showed that 5hmC modifications upon Lsh loss were not a direct consequence of changes in 5mC in these cells. Altogether, we identified the SNF2-like helicase LSH as a regulator of the hydroxymethylome at the genome-wide level.

## MATERIALS AND METHODS

### Cell Culture, transient transfection and Infection

Wild-type and Lsh KO MEFs (kindly received from Dr K. Muegge, (39)) were cultured in Dulbecco’s modified Eagle’s medium (DMEM) supplemented with 10% FBS and 100U/ml of penicillin/streptomycin. All cells were grown at 37°C in an atmosphere of 5% CO_2_. Wild-type, Lsh^off/off^ mouse embryonic stem cells (by Dr. Stancheva) were expanded on feeders using regular ES media (DMEM supplemented with 15% FBS, penicillin/streptomycin, non-essential amino acids, 1mM sodium pyruvate, 2mM L-glutamine and 100nM of β-mercaptoethanol) containing leukemia inhibitory factor (LIF). A modified knock-in strategy and allele design previously reported by Schnütgen et al (45) were employed to generate the Lsh^off/off^ ES cells by sequential targeted disruption of both *Lsh* alleles in E14 (129/Ola) ES cells. We first integrated by homologous recombination a reversible stop cassette (SA-GFP-Neo) flanked by a set of LoxP and Frt sites into the third intron of the *Lsh* gene. The integrated stop cassette is predicted to generate a null *Lsh* allele, which we named *Lsh^off^*, producing a chimeric protein containing 72 amino acids of the LSH N-terminus, which lacks nuclear localisation signal and any known function, fused to a GFP-Neomycin marker. The second Lsh allele was disrupted in one of the *Lsh^off/+^* ES cell lines by targeted integration of the same stop cassette, but this time carrying a hygromycin resistance marker. The successful integration of both stop cassettes was confirmed by Southern and Western blots. We will further refer to these *Lsh^off/off^* ES cells as Lsh KO ESCs.

Transient transfections were performed as described (46). Knock down in ES cells were designed accordingly to the authors’ instructions and performed in V6.5 ESCs (kindly received from Dr K. Koh, KU Leven) (47). Sequences used are KD1: CCGGCTAATCAGGGAGTTAAA, KD2: TCGAATGCTGCCCGAACTTAA, KD3: GGACACAGGATTAAGAATATG. Retrovirus production by HEK293GP cells and infection of target cells were performed as described (46). Infected cells were selected with 1.5μg/ml puromycin (Sigma).

### Halo Tag (HT) mammalian pulldown assay

HT mammalian pulldown assays were performed as previously described (10). Briefly, HEK293T cells were transfected with Halotag plasmid. Cells expressing HT-fusion proteins or HT-Ctrl were incubated in the mammalian lysis buffer supplemented with Protease Inhibitor cocktail (Promega) and RQ1 RNase-Free DNase (Promega) for 10min on ice.

The clarified lysate was incubated with HaloLink Resin (Promega) for 15min at 22°C with rotation. The resin was then washed with wash buffer and protein interactors were eluted with SDS elution buffer. Affinity purified complexes were then analyzed by nano-LC/MS/MS (MSBioworks) and Western blotting.

### Immunoprecipitation assays

Whole-cell extracts were prepared in IPH lysis buffer (50 mM Tris-HCl pH 8, 150 mM NaCl, 5mM EDTA, 0.5% NP40 supplemented with protease inhibitor cocktail (Roche). Immunoprecipitations were performed with anti-rabbit or anti-mouse IgG Dynabeads^®^ (Life Technologies) overnight at 4°C. The primary antibodies used in these experiments were directed against the following: control immunoglobulin G (IgG) (rabbit sc-2027 and mouse sc-2025; Santa Cruz), FLAG M2 (F3165; Sigma), LSH (NB100-278; Novus), TET1 (09-872; Millipore) and TET2 (R1086-4; Abiocode). The immunoprecipitated complexes were eluted in Laemmli buffer.

### GST pulldown assays

Full-length Lsh, Lsh CC (aa 1-226), Lsh DEXD (227-589) and Lsh CT (590-838) domains cloned in the pGEX4T-1 vector were kindly provided by Dr U. Ziebold (Cancer Research, Berlin, Germany) (48). Recombinant GST-fused proteins were expressed in and purified from Escherichia coli BL21, as previously described. We performed *in vitro* transcription/ translation using the TNT system (Promega) with pcDNA3 expression vectors for murine Flag-tagged Tet catalytic domains or full-length Halo-tagged human Tet proteins. GST-pulldown assays were performed as previously described (49).

### Dot blot

Genomic DNA was extracted with the DNeasy blood and tissue kit (Qiagen), denatured for 10 min at 95°C, chilled directly on ice and spotted on a nylon membrane (GE Healthcare Hybond-N^+^). After UV-induced fixation of target nucleic acids (2 x 200 000 μJ/cm^2^ of UV), the membrane was blocked in PBS BSA 1% and incubated with anti-5hmC (39769; Active Motif) or anti-ssDNA antibody (LS-C64821; LSBio).

### Analysis of global DNA 5mC and 5hmC levels by mass spectrometry (LC-MS/MS)

Analysis of global DNA 5mC and 5hmC levels by LC-MS/MS was carried out as described in Bachman *et al* (8). Briefly, 500 ng of genomic DNA was incubated with 5 U of DNA Degradase Plus (Zymo Research) at 37°C for 3 h. The resulting mixture of 2’-deoxynucleosides was analysed on a Triple Quad 6500 mass spectrometer (AB Sciex) fitted with an Infinity 1290 LC system (Agilent) and an Acquity UPLC HSS T3 column (Waters), using a gradient of water and acetonitrile with 0.1% formic acid. External calibration was performed using synthetic standards, and for accurate quantification, all samples and standards were spiked with isotopically labeled nucleosides. 5mC and 5hmC levels are expressed as a percentage of total cytosines (C, 5mC and 5hmC).

### hMeDIP

1μg of genomic DNA was diluted in ultra-pure water to 35 ng/μL and then sonicated in cold water with a Bioruptor sonicator (Diagenode) to obtain fragments averaging 300 bp in size. The fragmented DNA was used in combination with the hydroxymethyl collector (Active Motif) following the manufacturer’s protocol. Briefly, a glucose moiety that contains a reactive azide group was enzymatically linked to hydroxymethylcytosine in DNA, creating glucosyl-hydroxymethylcytosine. Next, a biotin conjugate was chemically attached to the modified glucose via a “click reaction”, and magnetic streptavidin beads were used to capture the biotinylated-hmC DNA fragments. After extensive washing steps and chemical elution, the hydroxymethylated DNA fragments released from the beads were used in sequencing experiments.

### Library preparation, deep sequencing workflow and data analyses

The library preparation was performed using the TruSeq ChIP Sample Prep Kit (Illumina). Briefly, double stranded DNA was subjected to 5’ and 3’ protruding ends repair and non-templated adenines were added to the 3’ ends of the blunted DNA fragments to allow ligation of multiplex Illumina’s adapters. DNA fragments were then size selected (300-500bp) in order to remove all non-ligated adapters. 18 cycles of PCR were done to amplify the library which was then quantified by fluorometry using the Qubit 2.0 and its integrity was assessed with 2100 bioanalyzer (Agilent) before being sequenced. 6pM of DNA library, spiked with 1% PhiX viral DNA, were clustered on cBot (Illumina) and sequencing was performed on a HiScanSQ module (Illumina). The BWA software was used to map sequencing reads to the mouse genome (NCBI Build 37/ UCSC mm9). Reads not uniquely mapped to the reference genome were discarded. Read density was computed by removing duplicate reads and by normalizing the local read count with respect to the total read count. Local enrichment in 5hmC was evaluated with the MACS software with a p-value cut-off < 10e-10. Differences between 5hmC levels were assessed for all annotated RefSeq transcripts (Gene Body) by first computing the normalized number of reads and then the fold change and absolute delta value between the two conditions. Genes were sorted according to their absolute fold change after discarding genes with a low absolute difference to avoid artefacts. 5hmC profiles around the TSS and within exons and introns were computed by first computing the local enrichment with the MACS software and then annotating the enrichment with the CEAS software. All statistics and plotting were done with the R statistical software. To obtain sequencing tracks, bedGraph files (genomeCoverageBed) were uploaded onto the IGV genome browser.

### Bioinformatic analysis

To identify the differentially hydroxymethylated regions, the genome was first structured in fixed windows (5000bp). The normalized 5hmC levels were then estimated by computing the FPKM for every window and each condition. The regions (windows of 5000bp) were then ranked based on their fold change and relative difference. Regions with an absolute fold change > 2 and absolute difference of at least 1 were selected for downstream analysis. Regions were related to genomic features using the VISTA Enhancer database (https://enhancer.lbl.gov), UCSC RefSeq and CpG islands annotations and by computing the genomic overlap between the region center and those features. The genomic regions between the transcription start site (TSS) and the transcription termination site (TTS) as defined as Gene Body, the 2kb regions upstream the TSS was defined as the promoter. Regions ambiguously overlapping multiple features were associated with multiple categories. For the metagene analysis all the RefSeq genes were used to compute the relative average 5hmC signal inside the transcript (TSS to TTS).

Public databases have been used: Histones marks ChIP-seq (GSE90893). RNA-seq data from ES cells GSM1581307 was used. Transcriptomic analyses for MEFs Lsh KO are downloaded from E-MEXP-2383 and methylation array from E-MEXP-2385(43). hMeDIP-seq on WT and Lsh KO MEFs and ES cells were uploaded under the GSE110129 number (token number: sfgrcoswrtybxej).

For the comparative analysis, ChIP-seq of histones marks were downloaded from GSE90893 and aligned onto the mm9 genome. Occupancy of 7 histones marks and one histone variant on the DhMR were then visualized with the seqMINER software(50). Free clustering was done and the count in each category with specific histone profile was performed.

### Repetitive element analysis (Pseudogenome)

A pseudogenome was generated with mouse DNA repeats sequences from RepeatMasker. Reads from fastq were mapped on this pseudogenome, using bowtie allowing two mismatches and without keeping reads mapped to more than one site. Duplicated reads were removed using samtools, and total reads mapped to each DNA repeats were calculated using samtools. The total numbers of reads mapped to each repeat element were normalized to the number of reads sequenced for each sample. To assess the effective change between the Lsh KO and the WT cells, the log odds ratio and P-value using a Fisher exact test were computed for each repetitive element. The P-values were then corrected using the FDR correction method.

### RT-qPCR and Gene expression

Total RNA was extracted with the RNeasy Mini kit (Qiagen). After DNase I treatment (DNA-free DNase kit, Ambion), Superscript II reverse transcriptase (Invitrogen) were used to reverse-transcribe mRNAs to cDNAs. Gene expression levels were then evaluated by real-time PCR (LightCycler 480, Roche). Primers used are available upon request.

### Ingenuity software

Ingenuity IPA software was used to identify the top molecular and cellular functions. The genes and regions lists were loaded into the Ingenuity database and then core analyses were done, using default parameters.

## RESULTS

### LSH is a TET-interacting factor

To explore the mechanisms of action of TETs, we identified TET protein partners via an unbiased proteomic approach using HaloTag technology as previously described (51). This approach identified several transcription factors and epigenetic proteins co-immunoprecipitating with TET proteins, including previously described binding partner OGT (10), PARP (52) and PCNA (53). We also identified the SNF2-helicase like LSH as a TET1 and TET3 binding partner with this method (data not shown).

To further explore the interaction between TETs and LSH, we first confirmed by semi-endogenous and endogenous co-IPs their interaction. As shown in Figure 1A, when HEK293T cells were transfected with Flag-tagged TET1, TET2 or TET3 catalytic domain (CD) and the empty vector. We found by immunoprecipitation that all TET CDs to interact with endogenous LSH (right, top panel, lane 6-8). LSH was not detected in the immunoprecipitated complex after transfection of cells with the Flag-empty plasmid (right, top panel, lane 5). Controls showing that the protein of interest was correctly immunoprecipitated (right, bottom panel) and that all proteins were expressed in cell lysates (left panel) are shown. These results were further confirmed in HEK293T cell lysates transfected with full-length Halo-tagged TET proteins (data not shown). The reverse IP was next performed (Figure S1A) confirming the interaction between LSH and TET proteins.

**Figure 1:**
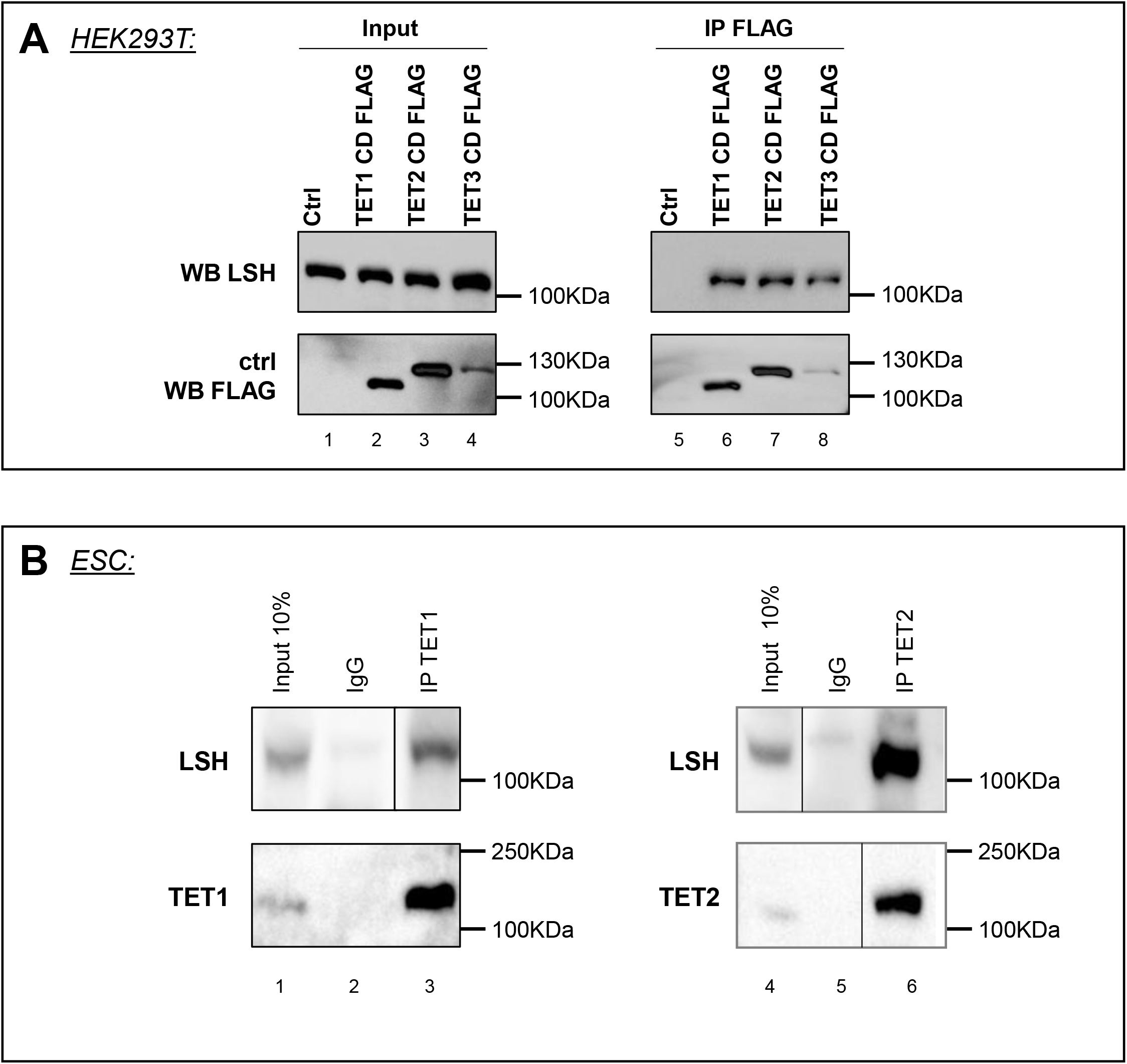
LSH is a TET interacting partner. (A) Halo pulldowns were performed from HEK293T cells overexpressing indicated Halo-tagged Tet protein. Spectral counts for LSH, SIN3A and OGT interacting proteins are indicated for biological replicates. As previously reported, TET1, TET3 but not TET2, shows interaction with SIN3A (75–77) and OGT interacts with all TETs, most abundantly with TET2 and TET3 (10). LSH interacts with TET1 and TET3. EEF1D (Isoform 1 of Elongation factor 1-delta) is a negative control. (B) Following transfection of the indicated FLAG-tagged constructs in HEK293T, cell extracts were precipitated with the anti-FLAG antibody and the presence of endogenous LSH was detected with anti-LSH antibody (right panel). Endogenous LSH was used as an input loading control and overexpression of TET catalytic domains (CD) is shown (left panel). (C) Endogenous co-IPs in ES cells. Cell lysates were subjected to immunoprecipitation with an anti-Tet1 (left panel) or anti-Tet2 (right panel) antibody and subjected to western blot using anti-Lsh or the indicated antibodies as control. Inputs and IP controls are shown. Vertical lines indicate juxtaposition of lanes within the same blot, exposed for the same time.

To map the LSH domain interacting with TET proteins, we performed additional GST pulldown assays. As shown in Figure S1B TET proteins were found to interact *in vitro* via their catalytic domain with the coiled-coil domain (CC) of LSH which is known to interact with DMNT1 and DNMT3B (48, 54).

To further validate the interaction between TET and LSH, we performed endogenous co-IPs in mouse embryonic stem cells using antibodies specific for LSH (Figure 1B) and for TET1 (left panel) or TET2 (right panel). We did not assess for TET3/LSH interaction in ES cells because TET3 is expressed at low levels in this cell system. Our results indicated endogenous interactions in ESC between LSH and TET1, and between LSH and TET2. Our data indicate that LSH and TET proteins interact *in vitro* and *in vivo*.

### Lsh knock-out impairs the global hydroxymethylation levels in MEFs and ES cells

Previous studies have shown that Lsh is involved in the establishment and maintenance of 5mC (35, 36, 41). Since DNA methylation is the substrate of hydroxymethylation, we thus decided to test whether Lsh could also contribute to 5hmC. To explore the role of Lsh in 5hmC, we performed dot blot experiments and mass spectrometry (MS) analyses (Figure 2). We prepared genomic DNA samples from Lsh KO and WT ES cells and analyzed them by dot blot with anti-5hmC and anti-single stranded DNA antibodies as a control. The quantification of the images by ImageJ showed that the levels of 5hmC are lower in Lsh KO compare to WT ES (Figure 2A). We reanalyzed the same samples by MS and confirmed the lower levels (29%) of 5hmC in the absence of Lsh without affecting the levels of 5mC (Figure 2B). We validated these observations by showing that depletion of Lsh by short-hairpin RNA (shRNA) in ES also causes a 50% reduction (shRNA 1), 32% (shRNA 2) and 70% (shRNA 3) of the 5hmC levels, which is similar to the 40% reduction observed by dot blot in KO ES cells (Figure S2 panel A and C). We also observed that another mark of demethylation, the 5-formylcytosine (5fC), is also dramatically reduced by 15-fold (Figure S2 panel B). Surprisingly, we observed that the levels of 5mC were similar between Lsh KO and control ES at global level, suggesting that Lsh loss directly reduces 5hmC establishment and/or maintenance rather than by an influence on DNA methylation.

**Figure 2:**
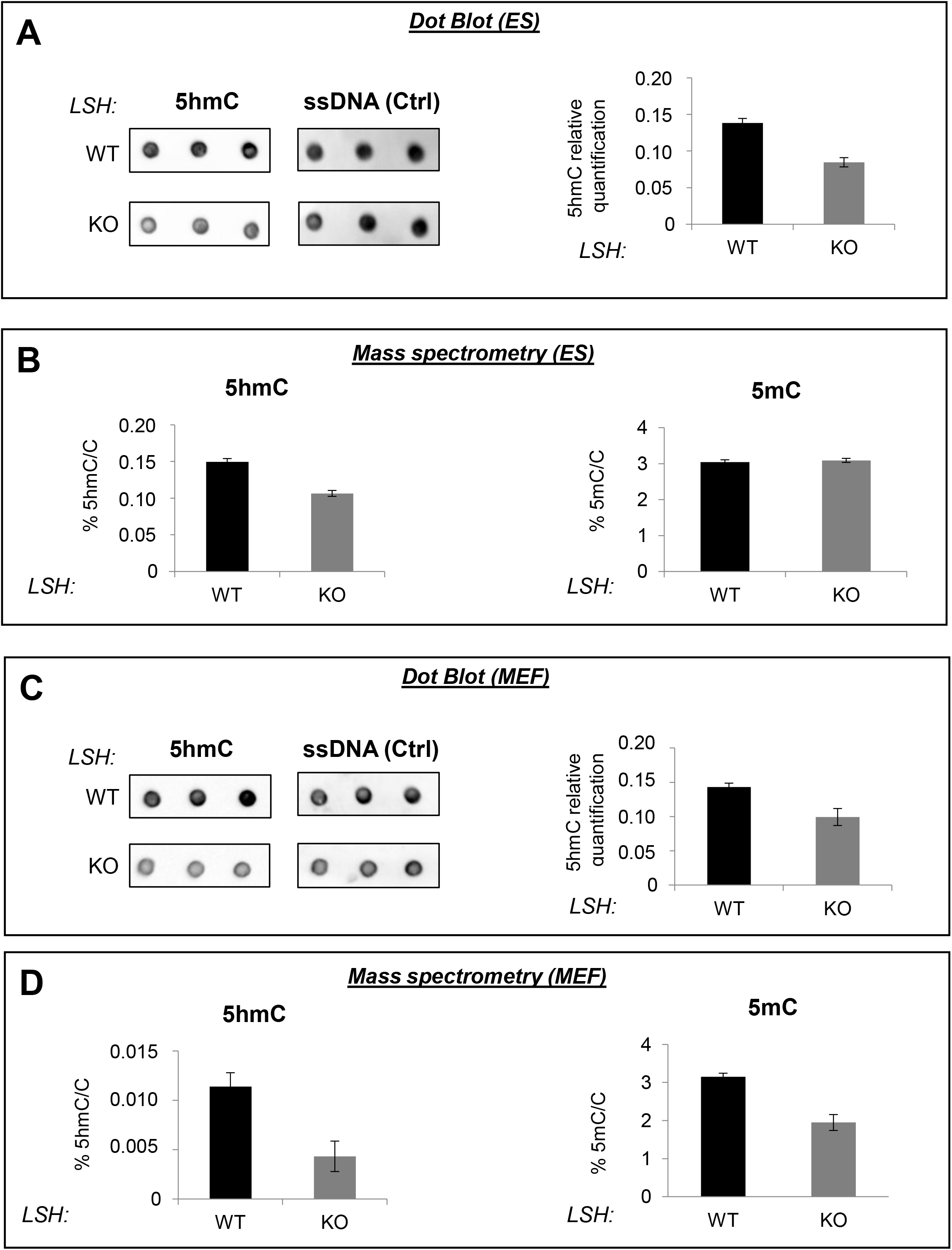
Lsh knock-out decreases 5hmC global level in ES and MEF cells. (A) Dot blot quantification of 5hmC global level in genomic DNA from wild-type and Lsh KO ES cells. ssDNA was used as loading control. Results were quantified using ImageJ. Error bars indicate s.d. of three biological replicates with a representative blot shown. (B) MS quantifications of 5hmC (left panel) and 5mC (right panel) in wild-type and Lsh KO ES. Error bars indicate s.d. of three biological replicates. (C) Dot blot quantification of 5hmC global level in genomic DNA from wild-type and Lsh KO MEF cells. ssDNA was used as loading control. Results were quantified using ImageJ. Error bars indicate s.d. of three biological replicates with a representative blot shown. (D) MS quantifications of 5hmC (left panel) and 5mC (right panel) in wild-type and Lsh KO MEFs. Error bars indicate s.d. of three biological replicates.

Our preliminary MS data showed that the levels of 5hmC are 10 times more abundant in ES than in MEFs cells. Nevertheless, we observed by dot blot that 5hmC levels were 1/3 lower in Lsh KO MEFs compared to WT MEFs (Figure 2C). Again, MS confirmed this result by showing a 63% reduction of 5hmC (Figure 2D). In contrast to what we detected in ES, we observed a reduction (by 40% at the genome-wide level) in the levels of 5mC by MS in the absence of Lsh as previously described (36, 43, 55).

Altogether, our results reveal that LSH maintain and/or establish 5hmC levels in ES and MEF cells. In ES cells, we could not detect significant changes in 5mC in the absence of Lsh suggesting that in this particular cell type Lsh could directly affect the function of Tet enzymes and 5hmC patterns. On the contrary, in MEFs cells, Lsh regulates both 5mC and 5hmC levels. Lsh may thus be involved in the regulation of 5hmC patterns in different ways according to the cell type.

### 5hmC changes at the genome-wide level in Lsh KO ESCs, mostly in gene bodies

To further explore the role of LSH in the distribution of 5hmC, we performed a genome-wide mapping of hydroxymethylation (hMeDIP-seq) and compared Lsh KO ESCs and MEFs Lsh KO to their wild-type counterparts (Figure 3 and Figure 4). Sequencing reads from Lsh KO and WT sample were aligned to the mouse mm9 genome. Using a window-based approach (5,000bp), we analyzed the read density along chromosomes and compared the density in Lsh KO and WT cells (cf Methods).

**Figure 3:**
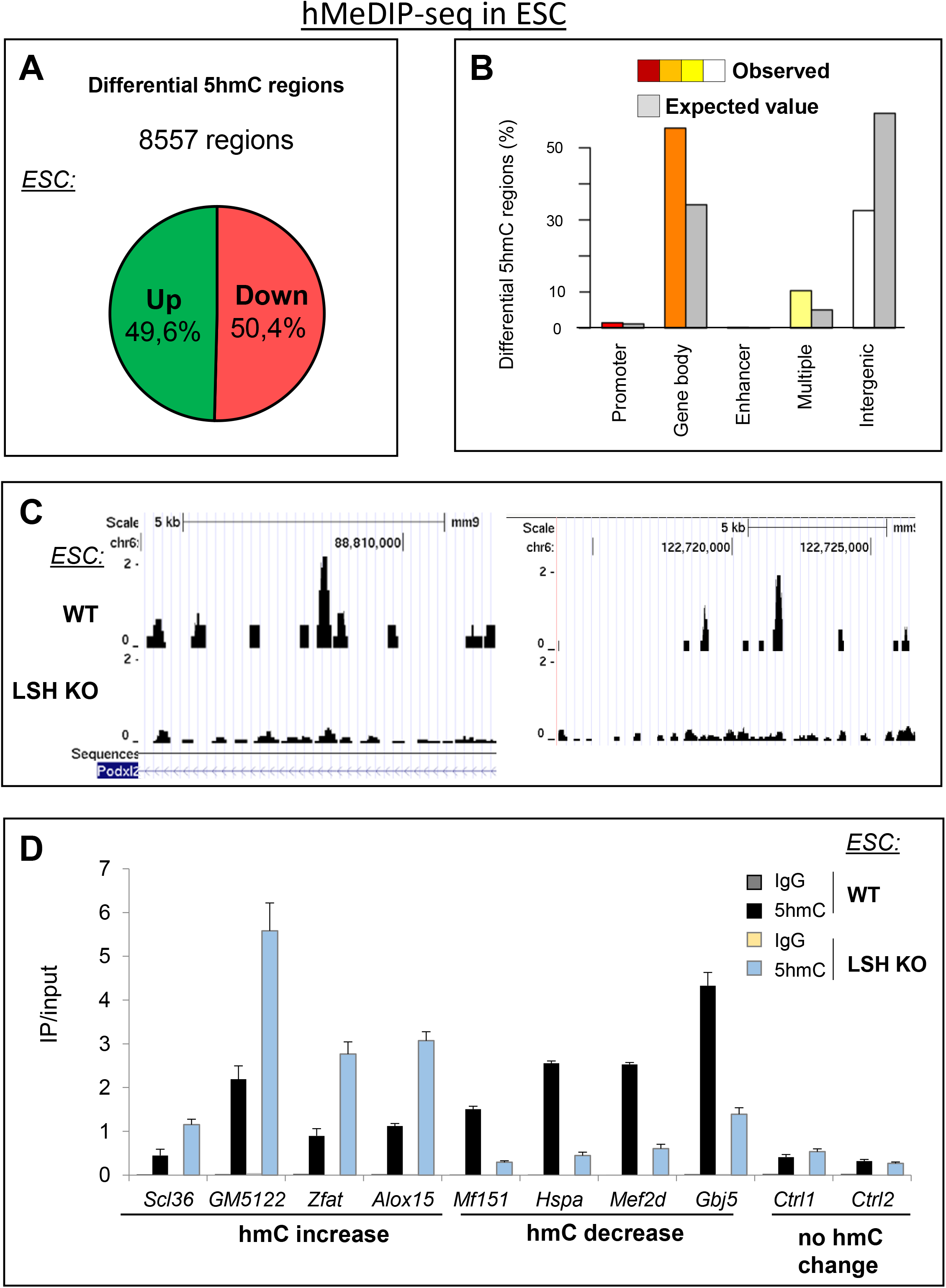
5hmC level is impaired in Lsh KO ES cells at specific genomic loci. (A) Pie chart showing the percentage of up and down 5hmC regions in ES cells upon Lsh loss (8557 regions in total). (B) Distribution of differentially hydroxymethylated regions on genomic features: promoters, gene body, enhancer, multiple or mixed features and intergenic. In color code are observed number whereas in grey are the expected numbers. (C) Representative UCSC Genome Browser plot from hMeDIP-seq data in ES cells (*Podxl2* gene). (D) An analyzed subset of differentially hydroxymethylated regions. qPCR analysis of hyper-hydroxymethylated genes (*Scl36, GM5122, Zfat and Alox15* genes) and on hypo-hydroxymethylated genes (*Mf151, Hspa, Mef2d and Gbj5*) after hMeDIP. IP/input represents real-time qPCR values normalized with respect to the input chromatin ± relative error of 3 independent experiments. Regions with no changes in 5hmC level are shown as negative controls (Ctrl1 & Ctrl2).

In Lsh KO ESCs, we identified 8557 windows (down 4312, up 4245, no statistical difference), corresponding to 5017 genes, showing differential 5hmC levels in ES cells, as shown for the representative gene *Podxl2* (Figure 3C). Half of these regions show gain and loss in 5hmC (Figure 3A). We found 2659 genes with gain of 5hmC and 2448 genes with loss of 5hmC (Supplemental Table 1). A deeper analysis of the distribution of these regions show enrichment at promoters (124/96 expected), gene bodies (4801/2940 expected), enhancers (13/5), multiple regions (i.e. region with at least two different genomic features, e.g. promoter and enhancer) (830/395 expected) and an underepresentation of intergenic regions (2789/5122 expected). These data indicate that regions with significant changes in 5hmC are mainly present in gene bodies and rarely occur at intergenic regions (Figure 3B). A functional analysis of hyper- and hypo-hydroxymethylated genes revealed significant over-representation of pathways associated with development, such as “tissue development”, “embryonic development” and “organismal development” (Figure S3D). We validated the changes in 5hmC profile by hMeDIP-qPCR. We confirmed that 5hmC was increased at *Scl36, GM5122, Zfat* and *Alox1*5 loci, while it was reduced at *Mf151, Hspa, Mef2d* and *Gbj5* (Figure 3D), as observed in hMeDIP-seq. No change of 5hmC was detected at two different control regions (Figure 3D).

5hmC changes often occur at gene bodies (>55%, Figure 3B). In contrast, significant changes in 5hmC rarely occur at promoters, but still in a higher than expected proportion (<3% of cases, Figure 3B), and at CpG islands (Figure S3A).

We investigate the 5hmC levels on the gene showing loss and gain of 5hmC upon LSH KO in the WT condition with Metagene analysis (Figure S3B). This analysis showed that both categories of genes have a higher level in 5hmC compared to the entire set of mouse genes. Nonetheless, no difference between the 5hmC level between hyper- and hypo-hydroxymethylated genes at the gene scale. A more detailed view of gene promoters showed a slight accumulation of 5hmC 1kb upstream to the TSS for hyper-hydroxymethylated genes (Figure S3C).

### 5hmC changes at the genome-wide level in Lsh KO MEFs

In Lsh KO MEFs, we identified 9002 tilling regions, corresponding to 3138 genes with differential hydroxymethylation. 77% of the regions (n= 6932) showed hypo-hydroxymethylation and 23% of the regions (n = 2070) harbor hyper-hydroxymethylation (Figure 4A) corresponding to 2376 and 1043 genes, respectively. As previously observed in ES cells (Figure 3B), these regions are mostly found in gene bodies (Figure 4B). Also, hydroxymethylation changes are rare at promoters (Figure 4B) and far from CpG islands (Figure S4A). No significant pattern is observed between genes hyper- and hypo-hydroxymethylated at TSS, gene body and TTS (Figure S4B). Changes in 5hmC level detected by high throughput sequencing were further confirmed by qPCR. *Bdnf, Btg4, Elfn2* and *Fam92* harbored an increased 5hmC whereas *Hp, Gdf5, Klf2* and *Sfi1* showed a decrease of 5hmC as in hMeDIP-seq (Figure 4C-D). No change of 5hmC was detected at control regions 1 and 2 (Figure 4D). Functional analysis of differentially hydroxymethylated genes showed enrichment in GO term “tissue development”, “tissue morphology” and “organismal development” (Figure S4C) highlighting again the global role of Lsh and the implication of (hydroxy)methylation in global development (41, 56, 57).

**Figure 4:**
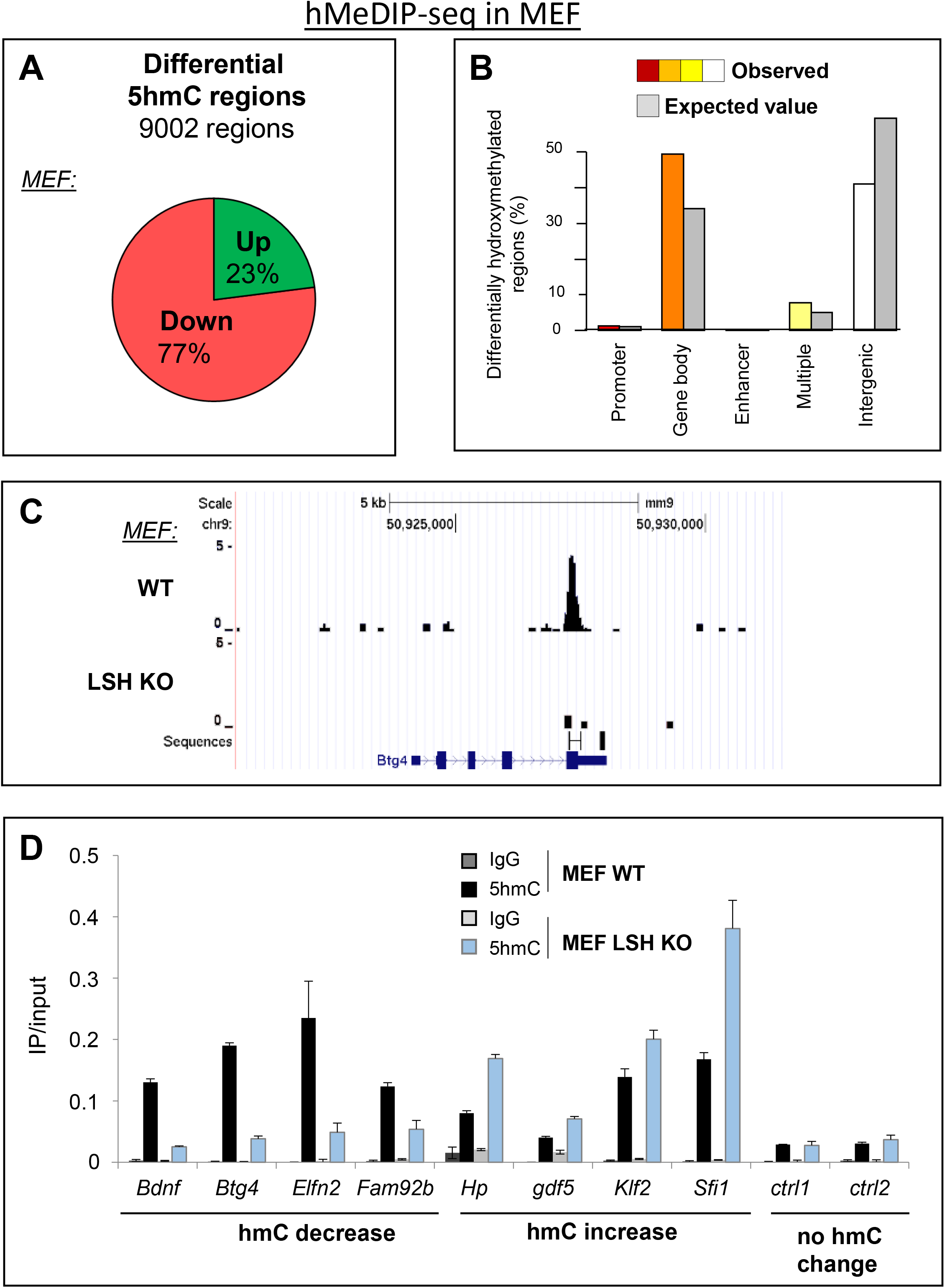
5hmC level is impaired in Lsh KO MEFs cells at specific genomic loci. (A) Pie chart showing the percentage of up and down 5hmC in MEF cells upon Lsh loss (9002 regions in total). (B) Distribution of differentially hydroxymethylated regions on genomic features: promoters (113), gene body (4504), enhancer (26), multiple or mixed features (652) and intergenic (3707). In color code are observed number whereas in grey are the expected numbers. (C) Representative UCSC Genome Browser plot from hMeDIP-seq data in MEF cells (*Btg4* gene). (D) An analyzed subset of differentially hydroxymethylated regions. qPCR analysis of hyper-hydroxymethylated genes (*Bdnf, Btg4, Elfn2 and Fam92b* genes) and on hypo-hydroxymethylated genes (*Hp, Gdf5, Klf2 and Sfi1* genes) after hMeDIP. IP/input represents real-time qPCR values normalized with respect to the input chromatin ± relative error of 3 independent experiments. Regions with no changes in 5hmC level are shown as negative controls (Ctrl1 & Ctrl2).

### Lsh regulates 5hmC at repetitive sequences

Lsh KO MEFs cells have been previously characterized by different groups (32, 55, 58, 59). A common finding is the global loss of 5mC, including at repeated minor satellite sequences. We directly accessed the level of 5hmC at major and minor satellite as well as repeated sequences such as LINE1 and SINE1 (Figure S5A). We observed by hMeDIP-qPCR a mild decrease in 5hmC at minor satellites and LINE1 elements in Lsh MEF KO cells, while a slightly decrease is detected at major satellite and a weak increase observe at SINE elements. We further expended the analysis to all DNA retreated sequences in the mice genome (Figure S5B). We mapped the reads on a synthetic pseudogenome containing all the repeated elements (Figure S5B) and identified several classes and families of repeated elements that exhibit changes in hydroxymethylation between Lsh KO MEFs and controls. In some cases, we observed loss of 5hmC at these repeated sequences. We next performed the same analysis in ES cells and found that major and minor satellites were hypo-hydroxymethylated. Thus, besides gene-regions, Lsh seems to influence hydroxymethylation of some repetitive sequences in ES and MEF cells.

### Relationship between 5hmC, mC and gene expression in MEF cells

In Lsh KO MEF cells, a common finding is the global loss of 5mC, including at repeated minor satellite sequences (34, 60). Myant and colleagues performed a genome-wide description of the global demethylation on Lsh KO MEFs (43). We used this array (RefSeq promoter microarray) to investigate the role of Lsh on 5mC and 5hmC pattern.

We first compared regions harboring deregulated methylation and hydroxymethylation in MEFs cells. We found only 17% of overlap and the overlap was not statistically significant (Figure 5A), which suggested that the patterns of DMRs and DhMRs were different. For instance, 5hmC changes observed at specific sites (Figure 4D), were not associated with changes in 5mC (Figure 5B). These findings suggest that 5mC and 5hmC patterns are not directly related to each other in MEF cells.

**Figure 5:**
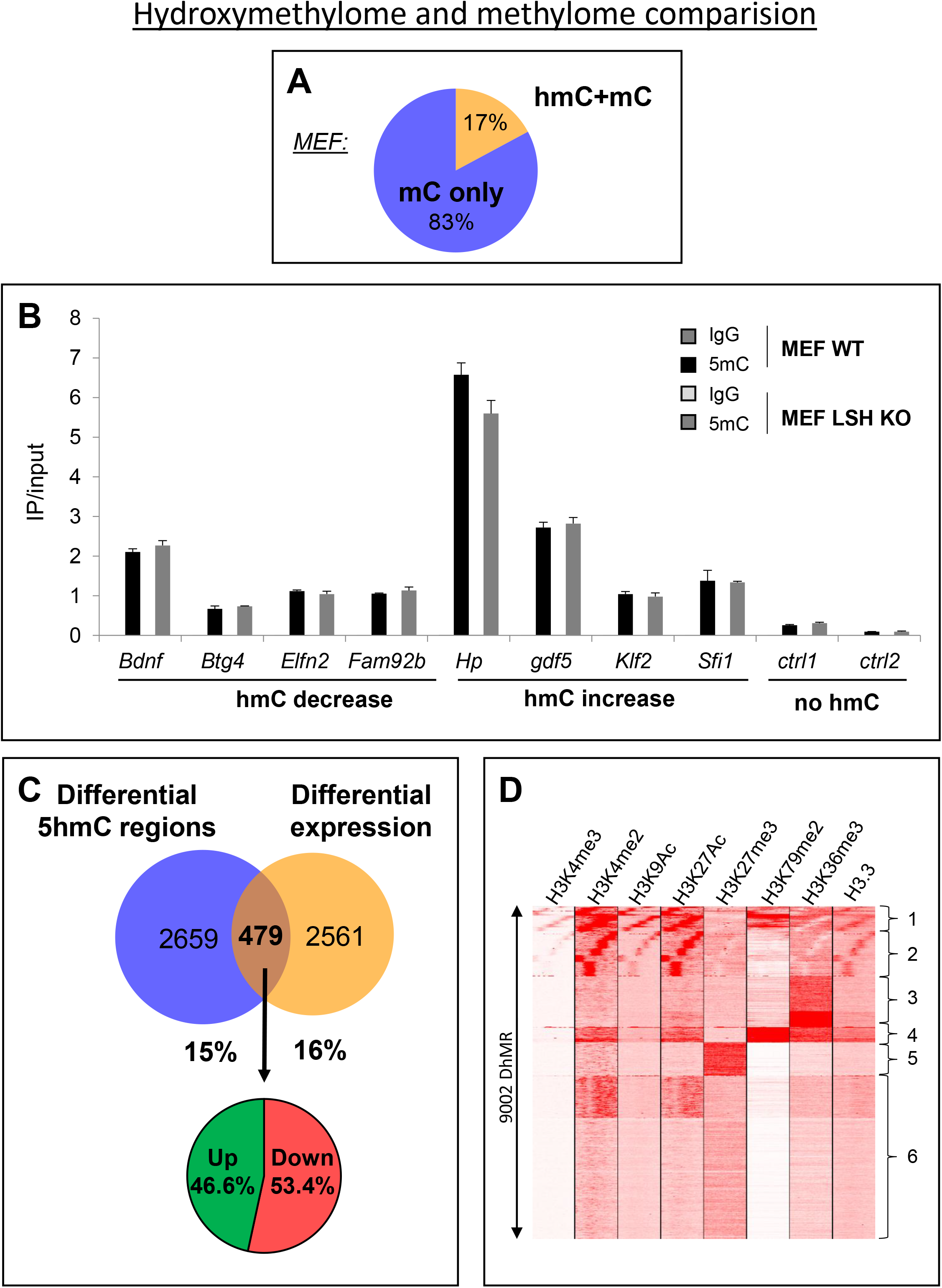
5hmC differences are not a direct consequence of 5mC modifications upon Lsh loss and are not always followed by direct gene expression change. (A) Pie chart representing the percentage of regions displaying changes in 5hmC level only, in 5mC level only or in 5hmC and 5mC level. Comparison was made between our 5hmC sequencing data and public available data from MeDIP followed by genome-wide tiling arrays in wild-type versus Lsh KO MEFs (43). (B) MeDIP qPCR experiments on target genes showing 5hmC increase, decrease or no 5hmC (control regions) in our high throughput sequencing data. Results are presented as percentages of Input ± relative error of 3 independent experiments. (C) Venn diagram showing overlap between the differentially hydroxymethylated genes identified by hMeDIP-seq (this study, see supp table 1) and differentially expressed genes (43). P-value overlap is 0.080300 showing no statistical relevance. Lower panel shows the mRNA up or down regulation of the 479 common genes. (D) Heatmap showing 7 histone modifications (H3K4me3, H3K4me2, H3K9Ac, H3K27Ac, H3K27me3, H3K79me2, H3K36me3) and one histone variant (H3.3) occupancy at the 9002 DhMRs defining 6 different sub-groups.

Myant and colleagues performed a global analysis of the transcriptionally misregulated genes in Lsh KO versus WT MEFs cells (43). We used these data to compare the regions showing differential 5hmC with expression data. We found (Figure 5C) that only 15% of genes that harbor DhMRs also have alteration of gene expression and 16% of genes that harbor differential expression have alteration of 5hmC level and the overlap was not significant. Also, no global expression changes were found between gene with deregulated 5hmC to all others (same result was obtained with hypo- and hyper-hydroxymethylated genes separately) (Figure S6). Thus, we confirmed that at the genome-wide level, changes in hydroxymethylation landscape around genes only partially affect their expression. Previous papers described also a poor overlap (~10%) between loss or gain of DNA methylation and gene expression changes (43).

In order to understand deeper the epigenetic status of the identified DhMRs, we next compared the 9002 DhMRs identified in MEFs cells to different histones marks that are characteristics of different chromatin environment or processes such as promoters (H3K4me3 and H3K9Ac), active promoters and enhancers (H3K4me2 and H3K27Ac), transcription (H3K36me3 and H3K79me2) or repressive compartments (H3K27me3) (61) (GSE90893) (Figure 5D). 21% of the DhMRs correlated with active histones marks: H3K27Ac, H3K4me2/3, H3K9Ac (group 1 and 2), 19% correlated with elongating marks as H3K36me3 and H3K79me2 (group 3 and 4) and only a few (10%) with repressive marks, H3K27me3 (group 5). For group 6, we found that half of the DhMRs do not harbor any examined marks. These data suggest the Lsh driven 5hmC is linked to different specific chromatin state and regions.

## DISCUSSION

LSH, a chromatin remodeler, was linked to DNA methylation and already known to interact with DNMTs (39, 40) and influence DNA methylation, in particular at repetitive sequences (33, 62). Here, we show that LSH is also able to interact with the DNA hydroxymethylases, the TET proteins. LSH binds the catalytic domain of TETs. On the other hand, TETs interact with the CC domain of LSH. CC domains are often involved in protein-protein interaction and this region of LSH is required for transcriptional silencing independently of its chromatin-remodeling activity (40). TETs and DNMTs both interact with the CC domain of LSH. Additional experiments are needed to answer the question of whether the binding of TETs and DNMTs to LSH is mutually exclusive.

The landscapes of 5hmC have been described for numerous cell type from human and mouse origin, but the molecular mechanisms involved in the deposition, maintenance and regulation of this epigenetic mark still remains unclear. Several studies have been performed to better understand the dynamics of 5hmC upon differentiation and cell growth changes (63–66). In addition, studies have been designed to identify the factors involved in 5hmC deposition and binding (11, 67, 68). Among these studies, one reported that LSH binds 5hmC containing DNAs *in vitro*, while LSH does not bind to the same oligonucleotide containing 5mC (9). It was also previously described that LSH can also induce the expression of TET enzymes (69). Then, LSH could modulate the 5hmC by increasing Tet mRNA level and/or influence their catalytic activity by interacting with 5hmC writers (9, 69). To decipher the exact contribution of each mechanism, producing a mutant form of Lsh that is unable to interact with TETs would be necessary.

In dot-blot, MS and deep sequencing experiments, we observed that the patterns of 5hmC were globally altered in ESCs and MEFs lacking LSH. These alterations in 5hmC are both gain and loss of 5hmC at specific genomic sites and repeated DNA sequences. Previous studies have shown a role of Lsh in 5mC in MEF cells. Comparison with published data (43) provides evidence that most changes of 5mC and 5hmC do not occur at the same genomic regions. Moreover, regions with 5mC changes do not correspond to changes of 5hmC. This is further consistent with a direct role of Lsh in establishment and/or maintenance of 5hmC pattern in ES and MEF cells. Unexpectedly, our results for the first time reveal a 5hmC regulation independently of 5mC patterns. The use of an auxin-inducible degron Lsh model could help to decipher precisely the dynamic of Lsh influence on ESCs and MEFs epigenomes.

Nevertheless, LSH seems to be a central regulator of DNA epigenome by influencing not only DNA methylation but also DNA hydroxymethylation. In particular, LSH can influence 5hmC by different mechanisms: (i) by interacting with TETs and potentially influencing their catalytic activity, (ii) by increasing Tets expression (70) and (iii) by binding to hydroxymethylated DNA (9).

The ES cells lacking LSH are viable and remain pluripotent indicating that changes in 5hmC and gene expression induced by Lsh KO affect, only modestly, the biology of ES cells. We postulate that changes in 5hmC levels upon Lsh KO are not sufficient to completely disorganize the transcriptional network of ES cells. 5hmC changes driven by Lsh KO are not associated with specific histone mark. However, 5hmC could generate a primed and more permissive state that could facilitate future transcriptional induction. It would be interesting to further investigate the dynamic of 5hmC and gene expression when Lsh KO ESCs differentiate into different lineages (ectoderm, mesoderm and endoderm). The functional analysis of genes affected by Lsh KO indicates that most of them are involved in development. Changes in 5hmC at specific genes in ES cells might be linked to changes in gene expression during specific steps of cell differentiation. Furthermore, it is also possible that LSH plays a broader function in genome organization beyond the local nucleosome remodeling activity. We observed that several families of DNA repeats exhibit changes in 5hmC upon Lsh KO in ES and MEF cells. It was also observed that the chromatin structure of repetitive sequences is impaired upon Lsh knock down (33, 71). These changes might contribute to the organization of specific chromosomal domains in the nucleus, linked to changes in epigenetic landscape and maybe gene expression in specific conditions. Further exploration is needed to uncover such mechanism.

LSH is located in a break point region frequently associated with leukemia (27) and a deletion in LSH gene is found in 57% of AML and 37% of ALL (72). Interestingly, 5hmC is often deregulated in hematological malignancies and TET2 is one of the most mutated genes in leukemia. In mice, loss of Tet2 increases the hematopoietic stem cell compartment and skews cell differentiation towards the myeloid compartment (73, 74). Both 5hmC levels and LSH are reduced in several solid tumors as nasopharyngeal carcinoma, breast or colon cancer (70). The link between LSH, TETs and 5hmC is not yet explored in hematopoietic diseases and further investigation could open new therapeutic possibilities.

In summary, we report here an interaction between TETs and LSH. Our genome-wide 5hmC profiles in MEFs and ES Lsh KO mouse cell lines suggest that LSH is an important regulator of DNA hydroxymethylation. Many efforts had been invested in studying the role of LSH in DNA methylation and little has been done on nucleosome positioning despite the classification of LSH as a full member of SNF2 ATPase remodeler family. Related to this function, we strongly suggest here a genome wide crosstalk between nucleosome positioning/chromatin organization and hydroxymethylation, which must be explored further.

## ACKNOWLEDGMENT

We thank Dr K. Muegge for the generous gift of MEFs Lsh WT and KO. We thank Dr M. Defrance for bioinformatical analyses and Dr C. Marchal for the technical help for repetitive region analysis. We thank Dr U. Ziebold for the generous gift of Lsh plasmids. We thank Dr B. Miotto for helpful discussions.

## AUTHORS CONTRIBUTIONS

MdD, RD and FF designed and coordinated the study. MdD, LC, EC run the experiments. MBi performed the bioinformatics analyses. MBa performed the MS experiments. CL and IS designed and generated the Lsh KO ESCs. MdD, LC, RD and FF wrote the manuscript. All the authors edited and approved the final manuscript.

## FUNDING

MdD and LC were supported by the F.N.R.S. MBi was supported by the Télévie. This work was funded by grants from the Fonds de la Recherche Scientifique and Télévie, as well as by grants from the IUAP P7/03 program, the Action de Recherche Concerté (AUWB-2010-2015 ULB-No 7), the Belgian “Foundation against Cancer,” the WB Health program, and the Fonds Gaston Ithier. Research in IS lab was supported by Cancer Research UK senior fellowship (C7215/A8983).

