## supplemental figures for "The chromatin remodeler LSH controls genome-wide cytosine hydroxymethylation"

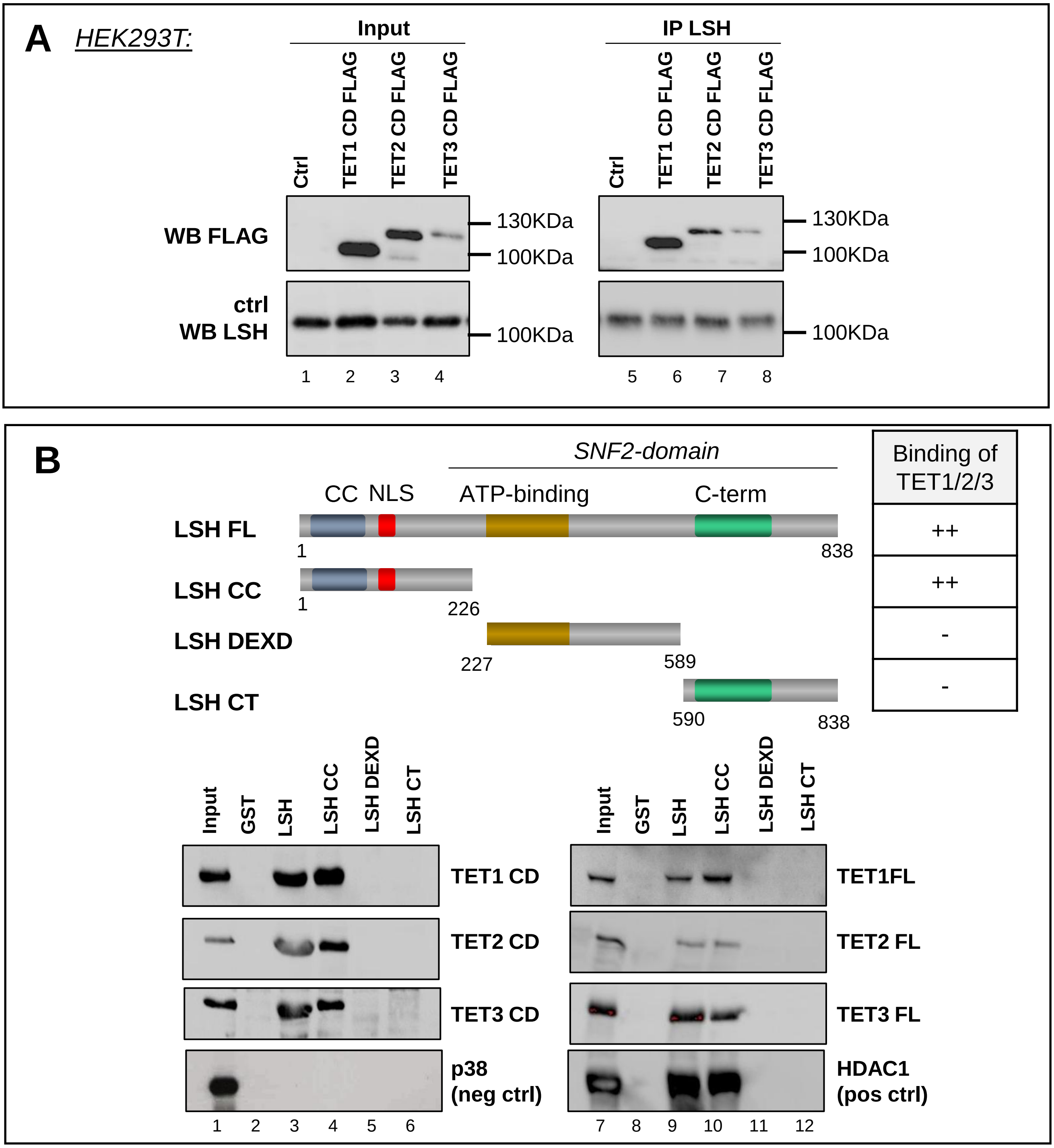


**Supplementary Figure 1 (related to Figure 1B). LSH associates with TET proteins.** (A) Immunoprecipitation with endogenous LSH antibody. Inputs in HEK293T are shown in left panel; upper panel is hybridized with FLAG antibody revealing catalytic domains of TET proteins and the lower panel with LSH antibody. Right part of the panel shows result of co-immunoprecipitation with LSH antibody revealing interaction of LSH with TET1/2/3 catalytic domain. (B) Upper panel: Schematic representation of the LSH protein, with its known domains highlighted (Von Eyss et al., 2012). Also shown are the different protein parts that were fused to GST and tested for binding to TET proteins. Lower panels: The indicated GST fusions were tested in GST pull-down experiments using IVT TET catalytic domains (left panel) or full-length TET (right panel). GST-HDAC-1 and GST-p38 were used as positive and negative controls, respectively.

**de Dieuleveult et *al.*, Supplementary Figure 1**

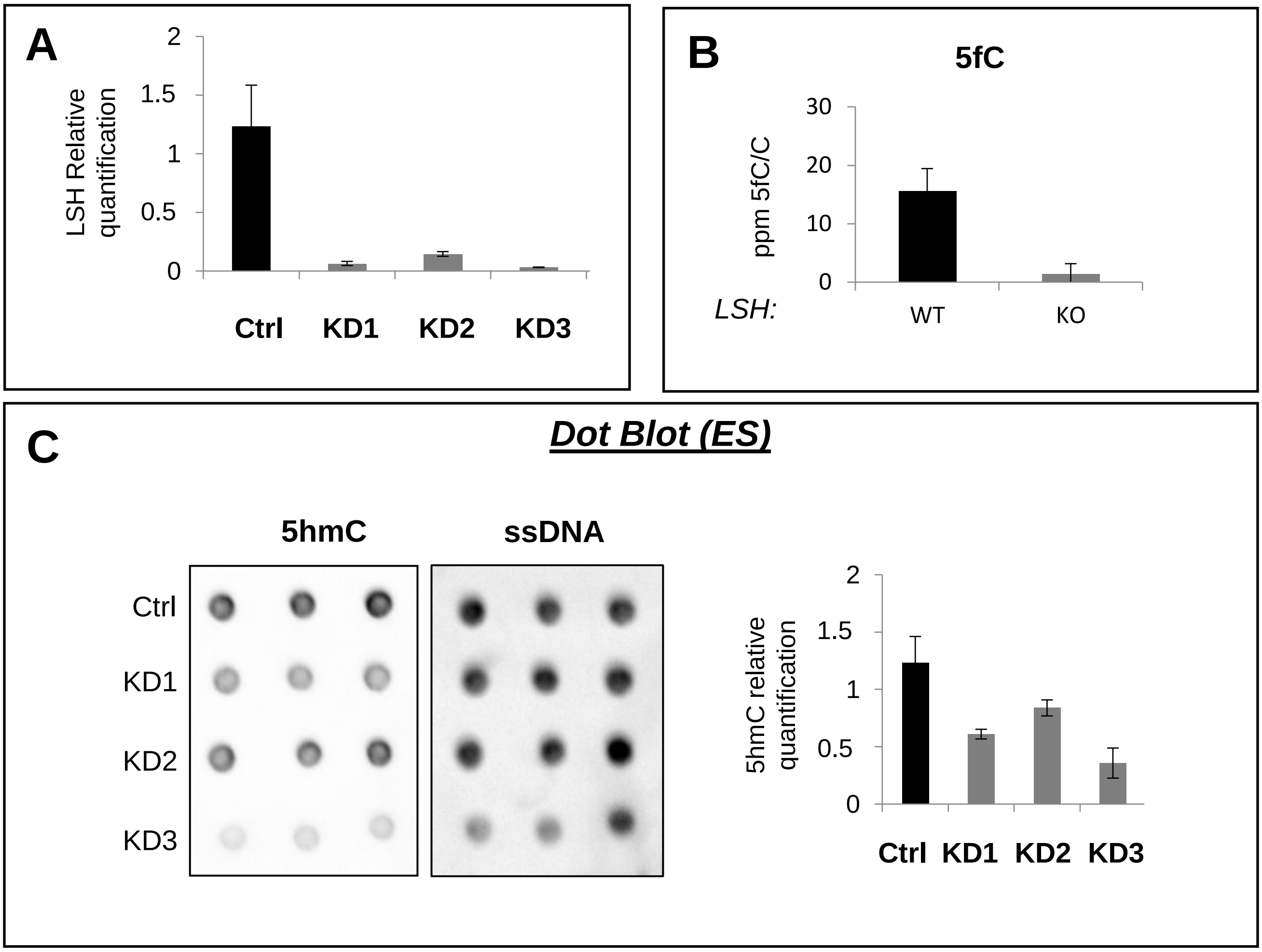


**Supplementary Figure 2 (related to Figure 2): Lsh knock-down decreases 5hmC global level in ES cells.** (A) Relative quantification of Lsh RNA level related to Supplemental Figure 2D normalized to GAPDH control. (B) 5fC relative quantification in Lsh WT and KO cells by mass spectrometry (C) Dot blot quantification of 5hmC global level in genomic DNA from wild-type and three different Lsh knock down ES cells (KD1, 2, 3). ssDNA was used as loading control. Results were quantified using ImageJ. Error bars indicate s.d. of three technical replicates with a representative blot shown.

### hMeDIP-seq in ES cells

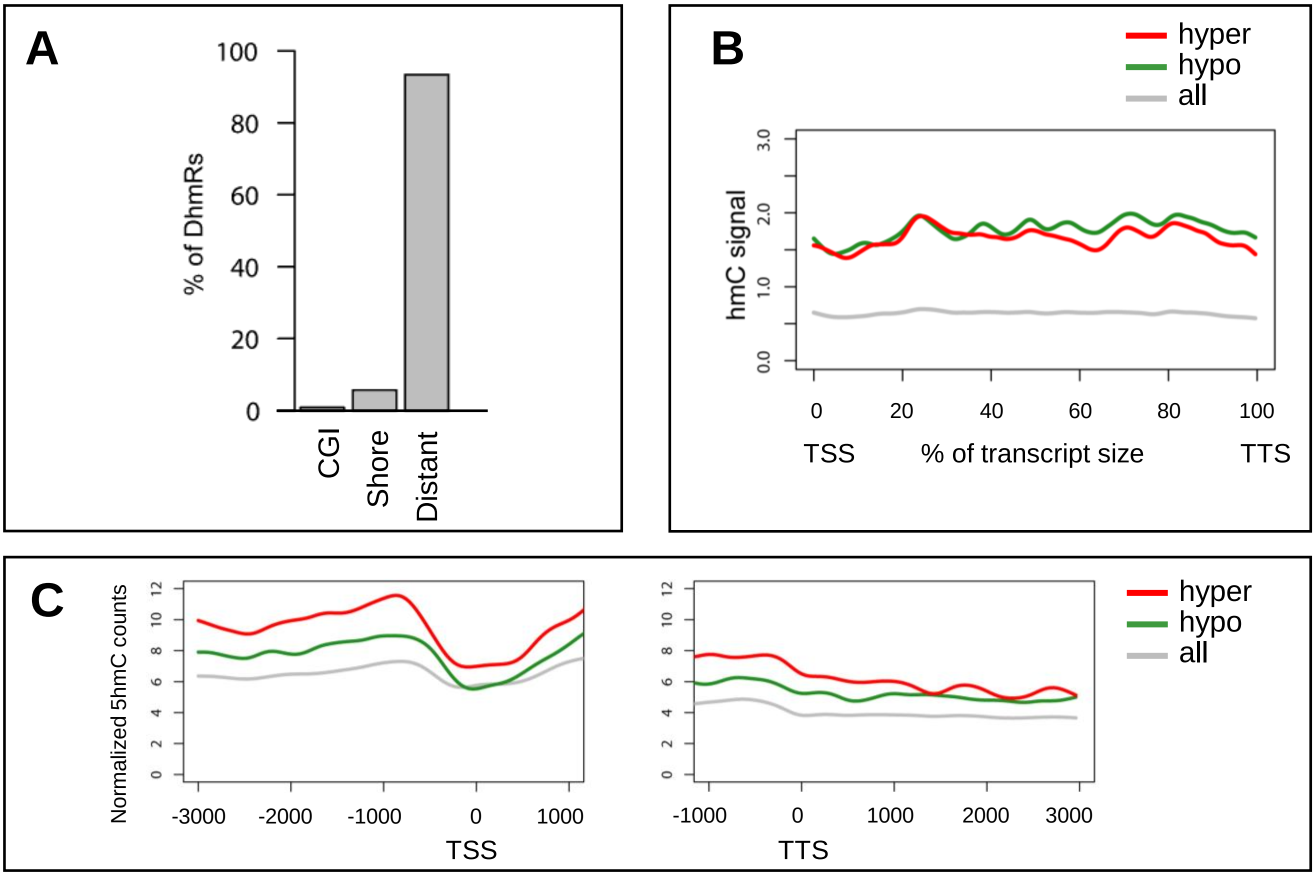


D

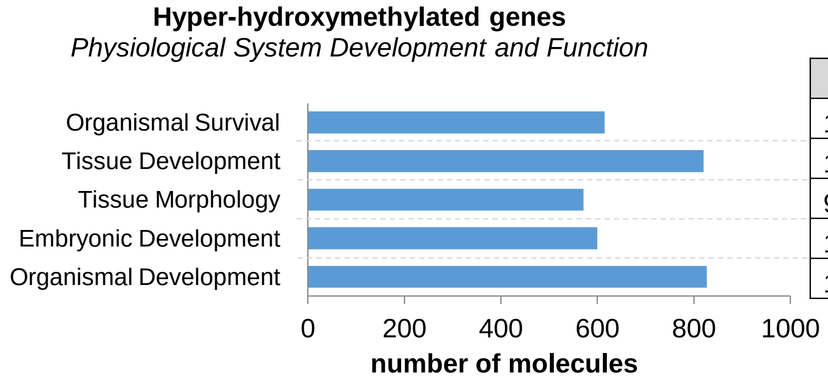


| p-value range | |
|---|---|
| 1.65E-10 | 7.67E-39 |
| 1.31E-04 | 2.86E-24 |
| 9.54E-05 | 2.00E-18 |
| 1.31E-04 | 2.05E-17 |
| 1.31E-04 | 2.05E-17 |

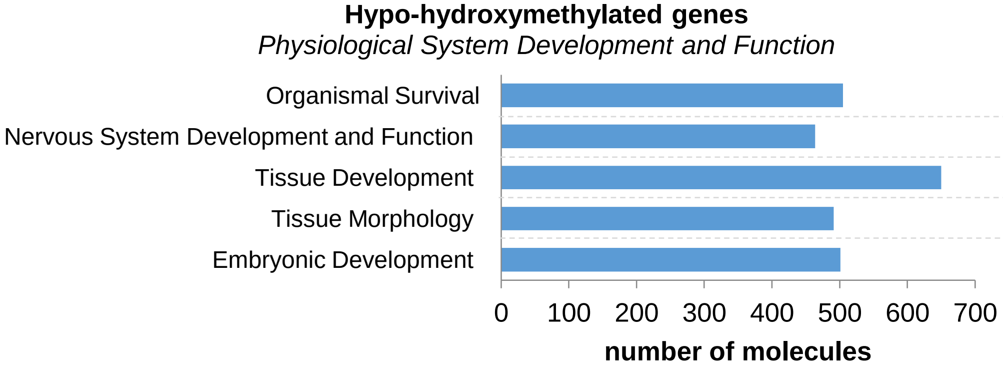


| p-value range | |
|---|---|
| 2.87E-04 | 1.51E-17 |
| 6.69E-04 | 7.39E-16 |
| 6.66E-04 | 7.39E-16 |
| 5.50E-04 | 6.63E-14 |
| 6.59E-04 | 4.26E-10 |

**de Dieuleveult et *al.*, Supplementary Figure 3**

**Supplementary Figure 3 (related to Figure 3): Genome wide DhMRs distribution and contribution to genomic landscape changes in ES cells.** (A) Percentage of differential hydroxymethylated regions (8557 DhMRs) versus genomic location: CpG island (CGI), close to CpG island (Shores) and distant to CpG islands (Distant) Expected values: CGI: 0.03326491, Shores: 0.0822699, Distant: 0.8844652.(B) Metagene profiles of 5hmC from transcription start site (TSS) to transcription termination site (TTS). All genes are scaled to fit the metagene profile (grey line), 5hmC profiles of all hyper hydroxymethylated genes are pooled altogether (red line) and 5hmC profiles of all hypo hydroxymethylated genes are polled altogether (green line).(C) GO ontology on 2659 genes with gain of 5hmC (upper panel) and 2448 genes with loss of 5hmC (lower panel). P-value range given by the Ingenuity software is indicated on right.

### hMeDIP-seq in MEF

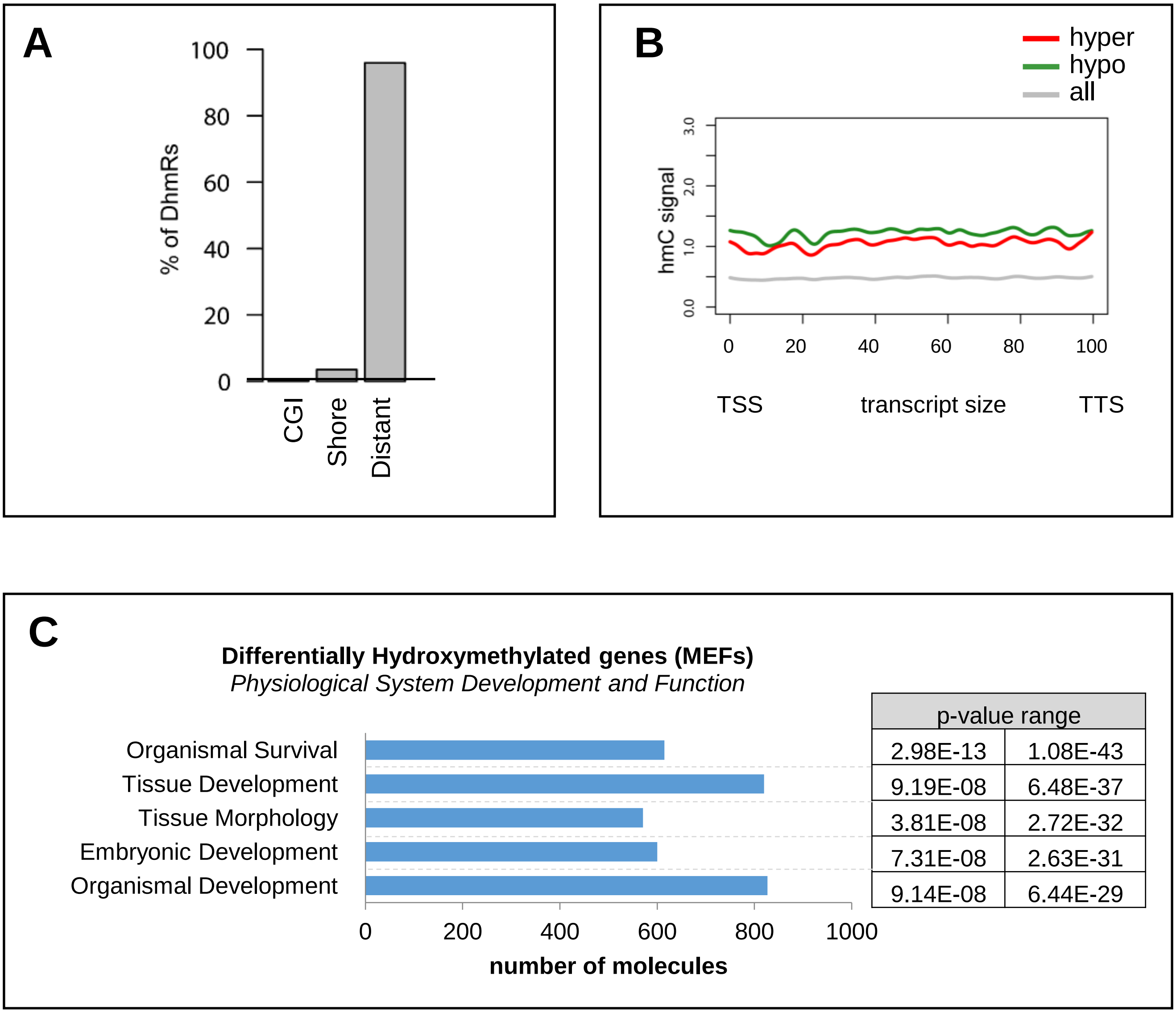


**Supplementary Figure 4 (related to Figure 4): Genome wide DhMRs distribution and contribution to genomic landscape changes in MEFs cells.** (A) Percentage of differential hydroxymethylated regions (9002 DhMRs) versus genomic location: CpG island (CGI), close to CpG island (Shores) and distant to CpG islands (Distant) Expected values: CGI: 0.03326491, Shores: 0.0822699, Distant: 0.8844652. (B) Metagene profiles of 5hmC from transcription start site (TSS) to transcription termination site (TTS). All genes are scaled to fit the metagene profile (grey line), 5hmC profiles of all hyper hydroxymethylated genes are polled altogether (red line) and 5hmC profiles of all hypo hydroxymethylated genes are polled altogether (green line). (C) GO ontology on 3419 DhmGe (genes with gain or loss of 5hmC). P-value range given by the Ingenuity software is indicated on right.

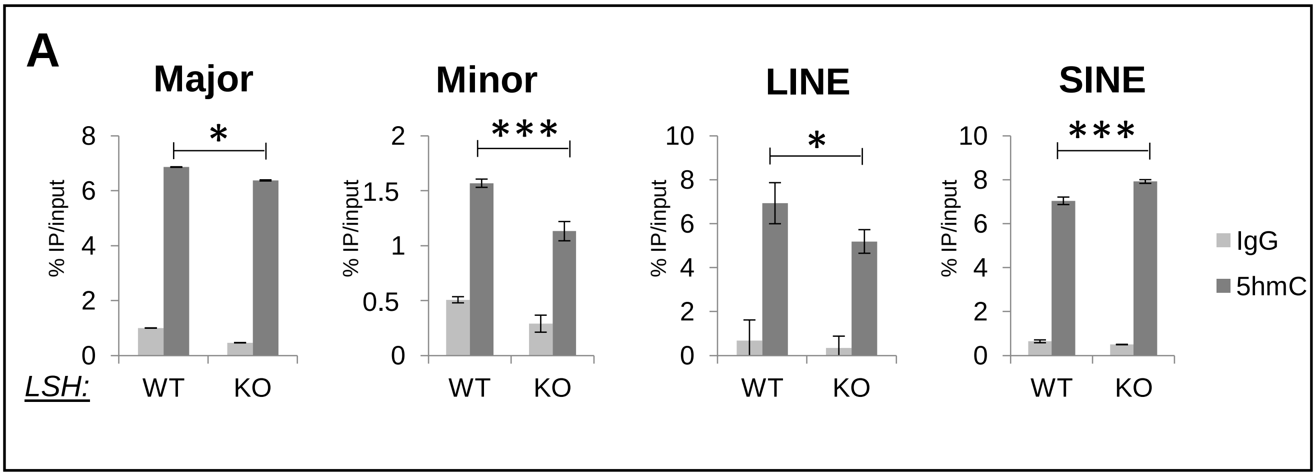


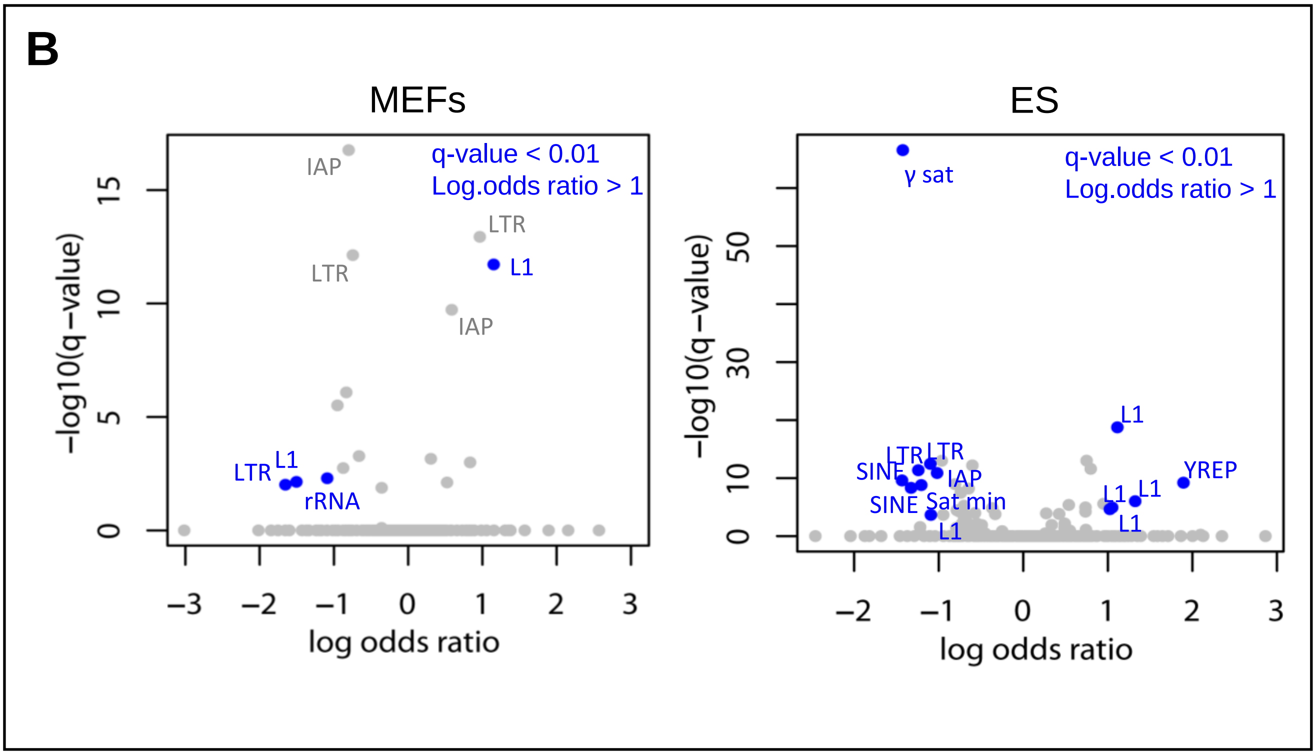


**Supplemental Figure 5: 5hmC is impaired at some repetitive sequences.** (A) qPCR following genome-wide hydroxymethylated DNA immunoprecipitation at the respective control with IgG on major satellites (Major), minor satellites (Minor), long interspersed elements (LINE) and short interspersed elements (SINE). Results are presented as percentages of input ± relative error of three independent experiments. (B) Scatter plot showing significance of the misregulation (-log10(q-value)) versus hydroxymethylation changes (log(odds ratio)) in WT and in KO MEFs cells (left panel) and in WT and in KO ES cells (right panel). Hydroxymethylation changes have been calculated by counting sequencing reads on pseudogenome analysis. Repetitive elements showing a q-value < 0.01 and a Log.odds ratio > 1 are indicated in blue.

**de Dieuleveult et *al.*, Supplementary Figure 5**

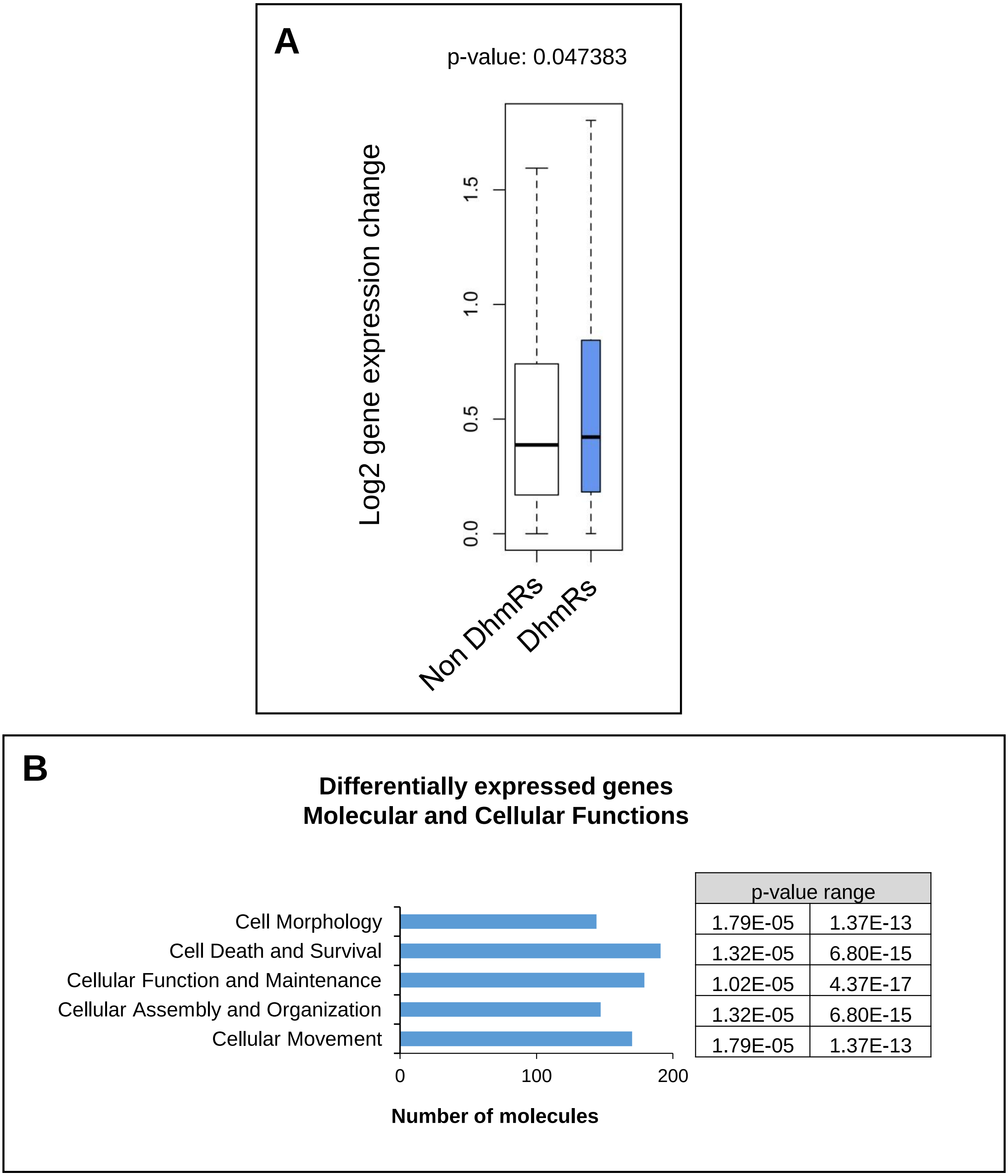


**Supplemental Figure 6: Analysis of the differentially hydroxymethylated genes.** (A) Histogram showing the average of all gene expression (microarray data, Myant et al.) compared to the average of DhMGe (hypo/hyper hydroxymethylated genes) expression in MEFs cells. T-test shows a p-value of 0.047383. (B) GO ontology on 479 genes with differential 5hmC level and differential expression. P-value range given by the Ingenuity software is indicated on right.

**de Dieuleveult et *al.*, Supplementary Figure 6**
